# Differential activation of P-TEFb complexes in the development of cardiomyocyte hypertrophy following activation of distinct GPCRs

**DOI:** 10.1101/2020.02.10.943209

**Authors:** Ryan D. Martin, Yalin Sun, Sarah MacKinnon, Luca Cuccia, Viviane Pagé, Terence E. Hébert, Jason C. Tanny

## Abstract

Pathological cardiac hypertrophy is driven by neurohormonal activation of specific G protein-coupled receptors (GPCRs) in cardiomyocytes and is accompanied by large-scale changes in cardiomyocyte gene expression. These transcriptional changes require activity of positive transcription elongation factor b (P-TEFb), which is recruited to target genes by the bromodomain protein Brd4 or the *S*uper *E*longation *C*omplex (SEC). Here we describe GPCR-specific regulation of these P-TEFb complexes and a novel mechanism for activating Brd4 in primary neonatal rat cardiomyocytes. The SEC was required for the hypertrophic response downstream of either the α_1_-adrenergic receptor (α_1_-AR) or the endothelin receptor (ETR). In contrast, Brd4 inhibition selectively impaired the α_1_-AR response. This was corroborated by the finding that activation of α_1_-AR, but not ETR, increased Brd4 occupancy at promoters and super enhancers of hypertrophic genes. Transcriptome analysis demonstrated that activation of both receptors initiated similar gene expression programs, but that Brd4 inhibition attenuated hypertrophic genes more robustly following α_1_-AR activation. Finally, we show that protein kinase A (PKA) is required for α_1_-AR stimulation of Brd4 chromatin occupancy. The differential role of the Brd4/P-TEFb complex in response to distinct GPCR pathways has potential clinical implications as therapies targeting this complex are currently being explored for heart failure.

## Introduction

The heart undergoes extensive remodelling in response to various mechanical and hormonal stressors during the progression to heart failure following myocardial infarction and/or sustained hypertension (1, 2). This includes hypertrophy of terminally differentiated cardiomyocytes in order to sustain cardiac output (3). While initially adaptive, prolonged cardiomyocyte hypertrophy leads to cardiomyocyte death, fibrosis and progression to chronic heart failure (4). Cardiomyocyte hypertrophy is initiated in part by neurohormonal activation of G protein-coupled receptors (GPCRs), such as the α_1_-adrenergic receptor (α_1_-AR), endothelin-1 receptor (ETR), and β-adrenergic (βAR) families (5, 6). Upon ligand binding, GPCRs activate heterotrimeric G proteins comprised of a Gα subunit and the obligate heterodimer Gβγ. Both the α_1_-AR and ETR canonically activate Gαq signalling, whereas the β-AR activates Gαs signalling. Cardiac-specific overexpression of Gαq and Gαs isoforms in mice leads to cardiomyopathy phenotypes, including cardiomyocyte hypertrophy (7, 8). The Gα isoforms elicit distinct signalling pathways involving calcium release and cyclic AMP (cAMP) formation respectively, which are capable of activating transcription factors and a gene expression program culminating in cardiomyocyte hypertrophy (9, 10). These pathological gene expression changes also require the coordinated interplay between dynamic alterations in chromatin structure, various master transcription factors and general transcriptional regulators, such as positive transcription elongation factor b (P-TEFb) (9, 11, 12).

P-TEFb, a heterodimer consisting of cyclin-dependent kinase 9 and cyclin T, positively regulates the release of RNA polymerase II (RNAPII) from a promoter proximal paused state into productive elongation. P-TEFb phosphorylates multiple proteins in the RNAPII elongation complex, including the C-terminal repeat domain of RNAPII itself, DRB sensitivity inducing factor (DSIF), and negative elongation factor (NELF) (13). In cardiomyocytes, P-TEFb activity is regulated by Gαq signalling as evidenced by cardiac-specific overexpression of Gαq in mice and ETR activation in primary neonatal rat cardiomyocytes (11). P-TEFb is a critical regulator for cardiomyocyte hypertrophy, with inhibition preventing the gene expression and cell size changes characteristic of cardiomyocyte hypertrophy (11). These transcriptional events require recruitment of the active P-TEFb complex to chromatin. P-TEFb recruitment is predominantly regulated through interactions with the bromodomain and extra-terminal (BET) protein Brd4 or through interactions with the super elongation complex (SEC) (14, 15).

Like other BET family members (Brd2, Brd3, and testis specific BrdT), Brd4 contains two N-terminal bromodomains, which bind to acetylated lysines on histone proteins leading to recruitment of Brd4 to chromatin, as well as an extra-terminal domain, which interacts with multiple transcriptional regulators (16). Brd4 and BrdT contain an additional domain that interacts with P-TEFb (17). The importance of Brd4 as a regulator of P-TEFb and transcription elongation in cardiomyocyte hypertrophy has been demonstrated using small molecule inhibitors of the BET bromodomain/acetyl lysine interaction such as JQ1 (18–21). JQ1 treatment reduced stress-induced gene expression and cardiomyocyte hypertrophy in primary culture models and in mice subjected to pressure overload via transverse aortic constriction (TAC), a potent inducer of cardiac hypertrophy *in vivo* (18, 21). Brd4 inhibition was also able to partially reverse pre-established signs of heart failure in a mouse model of myocardial infarction and pressure overload (20). These effects are correlated with the loss of Brd4 from super-enhancers and promoters of hypertrophic genes in cardiomyocytes, as well as reduced RNAPII elongation.

Multiple forms of SEC have been found in mammalian cells, comprised of P-TEFb, AF9, ENL, the three ELL family members (ELL1/2/3), EAF1/2, AFF1 and AFF4 (22). The SEC positively regulates release of RNAPII from promoter proximal-pausing to productive elongation (23). Aberrant targeting and activity of the SEC underlies development of various cancers and developmental diseases. For example, mixed lineage leukemia 1 (MLL) is fused to various SEC subunits in certain types of acute leukemias (24) and a germline Aff4 gain-of-function mutation leads to the developmental syndrome CHOPS (*C*ognitive impairment and coarse facies, *H*eart defects, *O*besity, *P*ulmonary involvement and *S*hort stature and skeletal dysplasia) (25). Although RNAPII promoter proximal pausing is dysregulated in cardiac hypertrophy, the role of the SEC in regulating the hypertrophic gene expression program in cardiomyocytes has not been investigated.

How diverse signaling pathways involved in cardiac remodelling cooperate to orchestrate the hypertrophic gene expression program *in vivo* remains poorly understood. Although neurohormonal signals induce similar hypertrophic responses in primary cardiomyocytes, distinct signalling pathways are initiated through activation of their cognate GPCRs (26). Such effects are generally attributed to differential G protein coupling, however receptor-specific differences in downstream effector protein activation of the same Gα subunit have also been demonstrated (27). The coordination between these signalling pathways, and the fact that each activates a unique combination of transcription factors, suggests that there may be differences in how they regulate gene expression. However, comparison of changes in gene expression have only been assessed for a limited repertoire of genes (28, 29). How different receptors alter global transcriptional regulation has not been systematically compared. Such differences may have important therapeutic implications for treating patients with heart disease stemming from varied clinical origins.

In this study we focused on the differential impact of cardiomyocyte GPCR signalling pathways on mechanisms regulating transcription. We investigated the role of P-TEFb and its interacting partners, Brd4 and SEC, in cardiomyocyte hypertrophy caused by activation of either of two GPCRs, the α_1_-AR or ETR. These receptors are canonically thought to elicit their hypertrophic responses through Gαq activation, with both receptors also able to activate additional Gα subunits which has not been thoroughly assessed. We found that P-TEFb activity and the SEC are required for cardiomyocyte hypertrophy induced by activation of either GPCR. However, only the α_1_-AR response was attenuated by Brd4 inhibition. Transcriptome analysis after Brd4 inhibition indicated attenuation of α_1_-AR upregulated genes that were enriched for pathways involved in the pathophysiology of cardiomyocyte hypertrophy. Brd4 chromatin occupancy at promoters and super enhancers of hypertrophic genes was specifically induced by α_1_-AR activation, an effect that was dependent on the activity of PKA. Lastly, we demonstrated that the hypertrophic response downstream of another receptor known to signal through PKA, the β-AR, was also attenuated by Brd4 inhibition. Our study suggests receptor-specific regulation of P-TEFb function and expands the currently known cellular repertoire of protein kinases capable of regulating Brd4 function. Further, our findings suggest that the clinical efficacy of BET inhibitors for heart failure may depend on patients’ specific neurohormonal signalling patterns.

## Results

### Evidence for receptor-specific P-TEFb regulation in cardiomyocyte hypertrophy

We first revisited the requirement of P-TEFb activity for the hypertrophic response in primary neonatal rat cardiomyocytes (NRCMs). Previous experiments used the ATP analog 5,6-dichlorobenzimidazone-1-β-D-ribofuranoside (DRB), a cyclin-dependent kinase inhibitor that affects Cdk9, to implicate P-TEFb activity in the hypertrophic response (11). To confirm these results, we repeated this experiment using iCdk9, a Cdk9 inhibitor that is ∼1000 times more potent and ∼100 times more selective than DRB (30). Following 24 h treatment, agonists for the ETR (endothelin-1; ET-1) or α_1_-AR (phenylephrine; PE) increased cardiomyocyte surface area by 35-40% relative to control, as assessed by analysis of α_2_-actinin immunostaining using high-content microscopy (Figures 1A and 1B). Simultaneous treatment with 0.2 µM iCdk9 completely abolished the increase in cardiomyocyte size elicited by either agonist, confirming a stringent requirement for P-TEFb activity in cardiomyocyte hypertrophy (Figures 1A and 1B).

**Figure 1.**
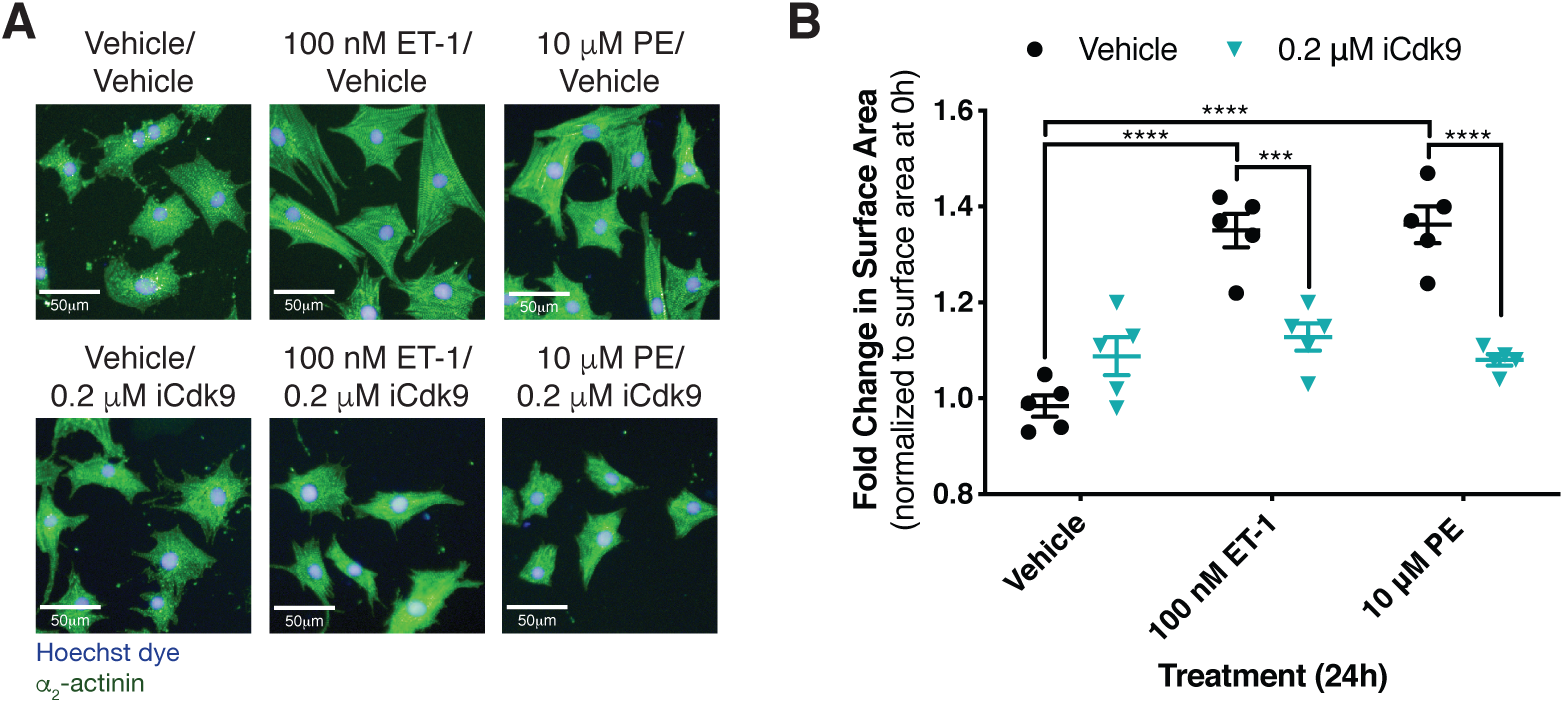
Inhibition of the P-TEFb kinase subunit Cdk9 prevents cardiomyocyte hypertrophy in response to α1-AR or ETR activation. **(A)** NRCMs were treated with PE or ET-1 for 24 h as indicated. Cardiomyocytes were stained with Hoechst dye and identified by staining for the cardiomyocyte-specific marker α_2_-actinin. **(B)** Fold change in cardiomyocyte surface area following 24 h treatment over the surface area of cardiomyocytes from the same biological replicate at 0 h. Data is presented as mean ± S.E.M with each point representing a biological replicate. Two-way ANOVA followed by post-hoc t-tests with Bonferroni correction was performed (***p<0.001, ****p<0.0001).

To further characterize P-TEFb function in the hypertrophic response, we assessed the roles of the SEC and Brd4, two key P-TEFb-interacting proteins (31). To perturb the P-TEFb/SEC interaction, we used KL-2, which prevents the interaction between the cyclin T component of P-TEFb and the SEC scaffolding subunit Aff4 (32). KL-2 treatment blocked the increase in cell size in response to both ETR and α_1_-AR activation, similar to the effect of Cdk9 inhibition (Figures 2A and 2B). To determine whether disruption of the P-TEFb/SEC interaction also affected the expression of hypertrophic genes, we monitored changes in mRNA levels for established hypertrophy marker genes Nppa, Nppb, and Serpine1 using RT-qPCR. Whereas mRNA levels for these genes were robustly increased in response to activation of either receptor, induction of Nppb and Nppa was blocked by KL-2 co-treatment (Figure 2C). Interestingly, Serpine1 induction was unaffected by KL-2 treatment indicating a gene-specific aspect to SEC function. To confirm the observed effect was due to disruption of SEC function, we next reduced Aff4 levels using siRNA and verified knockdown by RT-qPCR (Figure 2D). Similar to KL-2 treatment, knockdown of Aff4 blocked the increase in cell size following activation of either receptor (Figure 2E).

**Figure 2.**
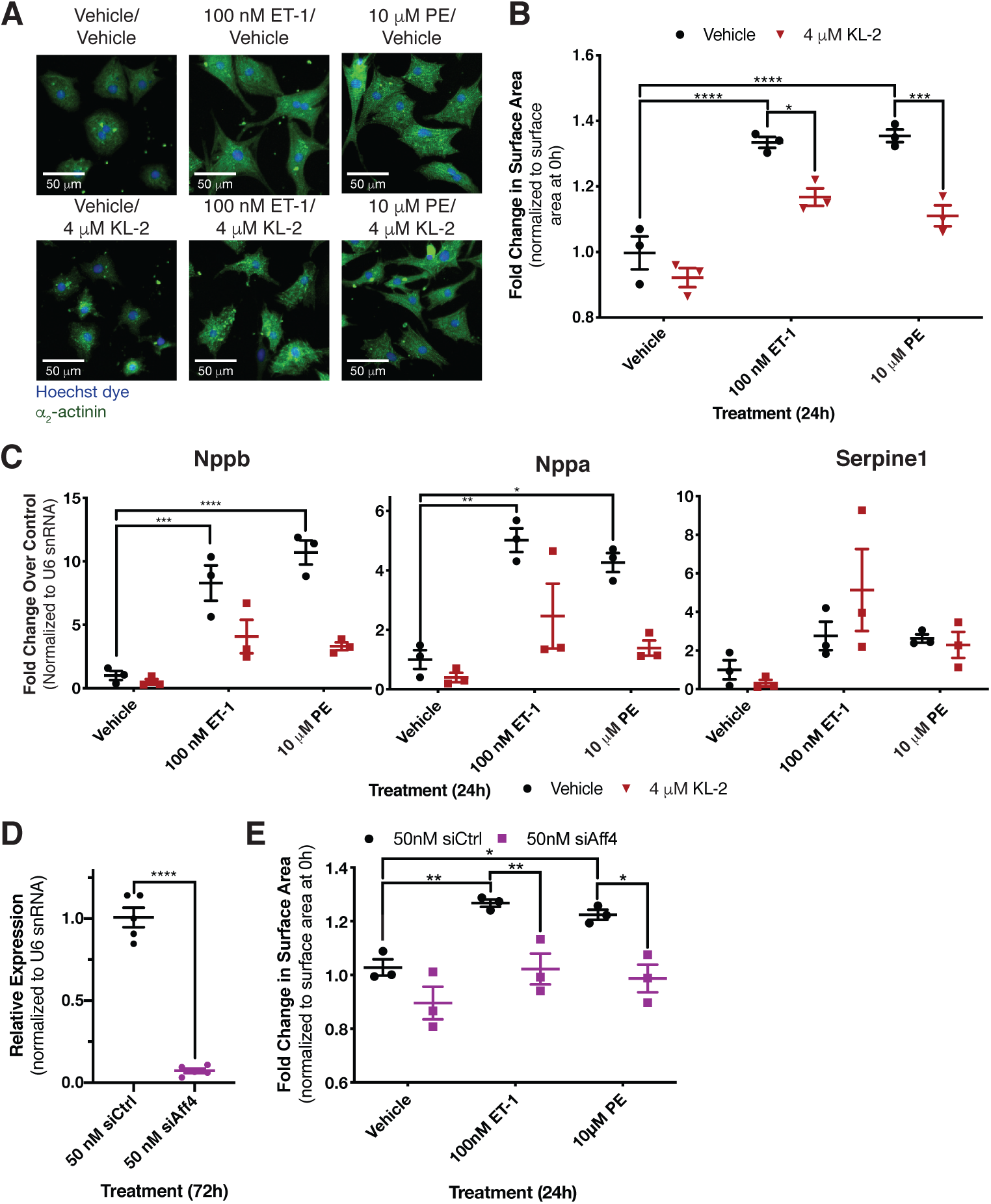
Disruption of SEC-P-TEFb interaction blocks the hypertrophic response following activation of either receptor. **(A)** NRCMs were treated for 24 h as indicated. Cardiomyocytes were stained with Hoechst dye and identified by staining of the cardiomyocyte-specific marker α_2_-actinin. **(B)** Fold change in surface area of identified cardiomyocytes over siRNA control-transfected cardiomyocytes from the same biological replicate at 0 h. Two-way ANOVA followed by post-hoc t-tests with Bonferroni correction was performed. **(C)** Expression of three genes previously identified as upregulated in hypertrophic cardiomyocytes, Nppb, Nppa and Serpine1, was determined by RT-qPCR. Two-way ANOVA followed by post-hoc t-tests with Bonferroni correction was performed. **(D)** Aff4 knockdown in cardiomyocytes 72 h after transfection with Aff4 targeted siRNA was validated by RT-qPCR. An unpaired t-test was performed. **(E)** Fold change in surface area of identified cardiomyocytes over the surface area of siRNA control-transfected cardiomyocytes from the same biological replicate at 0 h. Cardiomyocytes were transfected 72 h prior to treatment with 50 nM of the specified siRNA. Data is presented as mean ± S.E.M with each point representing a biological replicate. Two-way ANOVA followed by post-hoc t-tests with Bonferroni correction was performed (*p<0.05, **p<0.01, ***p<0.001, ****p<0.0001).

We next tested the role of Brd4 using the pan-BET small-molecule inhibitor JQ1. JQ1 targets the BET family bromodomains, acting to competitively inhibit their interaction with acetylated lysine residues (33). Interestingly, BET inhibition attenuated the hypertrophic response in a receptor-specific manner: the response to α_1_-AR activation was decreased, whereas there was no effect on ETR-mediated increase in cell size (Figures 3A and 3B). We also observed that JQ1 treatment more strongly reduced expression of hypertrophic marker genes in cells stimulated with the α_1_-AR agonist compared to cells stimulated with an ETR agonist (Figure 3C). One possible explanation for the receptor-specific effect of JQ1 is that the dose of ET-1 used to drive hypertrophy was sufficiently high to overcome JQ1 inhibition. To address this, we tested the effect of JQ1 on the ET-1-driven hypertrophic response over a wide range of ET-1 doses. The ET-1 response was insensitive to JQ1 at all doses tested (Figure 4). Thus, receptor-specific differences in JQ1 sensitivity likely reflect intrinsic differences in the respective GPCR signalling outcomes. These data suggest that the SEC is generally required for P-TEFb function in the hypertrophic response, whereas the P-TEFb/Brd4 complex mediates receptor-specific functions.

**Figure 3.**
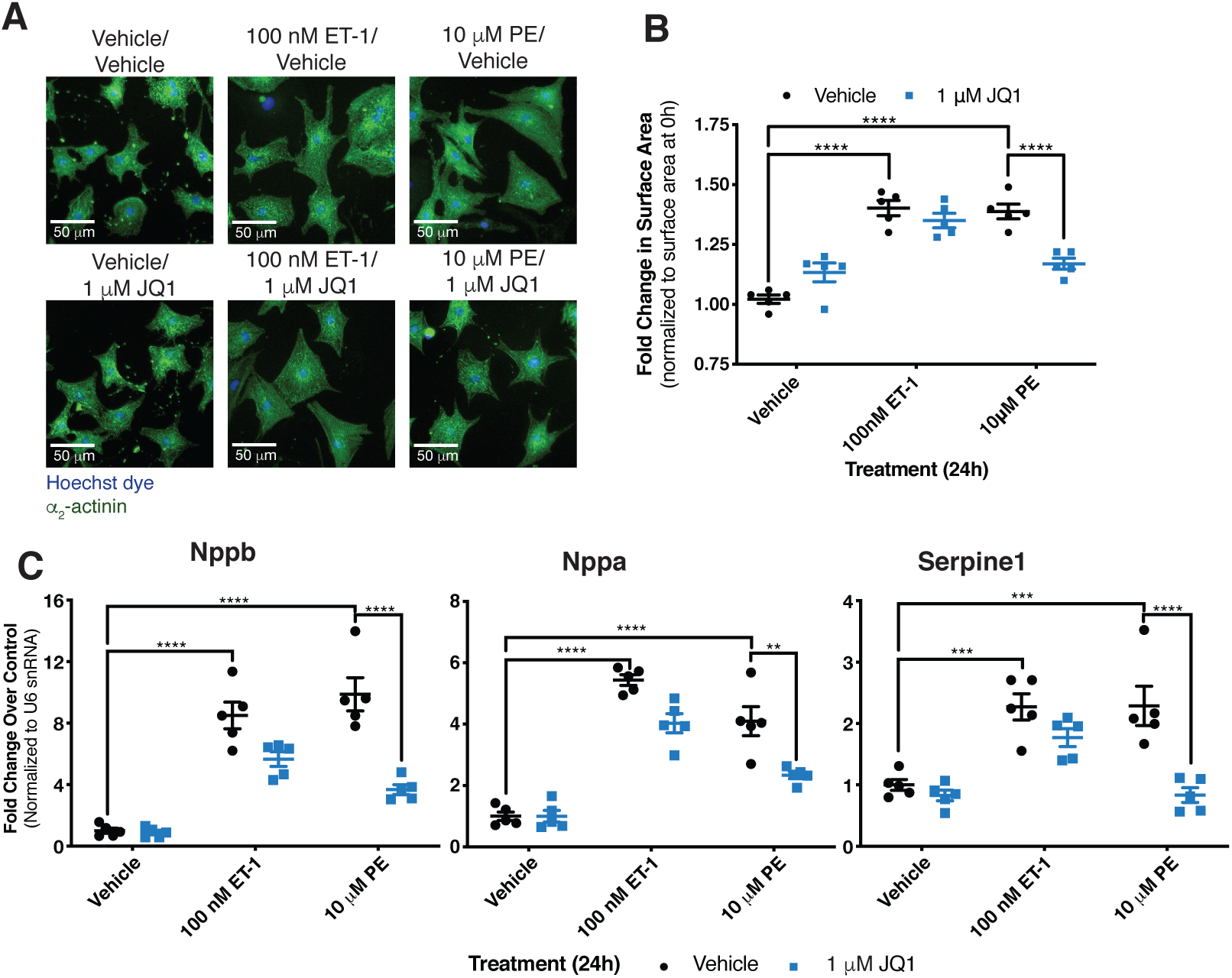
Effects of BET inhibitor JQ1 on cardiomyocyte hypertrophy are specific to the receptor driving the response. **(A)** NRCMs treated for 24 h were fixed and stained with Hoechst dye and identified by staining for the cardiomyocyte-specific marker α_2_-actinin. **(B)** Fold change in surface area of cardiomyocytes over surface area of cardiomyocytes at 0 h from the same biological replicate. Two-way ANOVA followed by post-hoc t-tests with Bonferroni correction was performed. **(C)** Expression of genes previously demonstrated to be upregulated in hypertrophic cardiomyocytes was determined by RT-qPCR. Data is presented as mean ± S.E.M with each point representing a biological replicate. Two-way ANOVA followed by post-hoc t-tests with Bonferroni correction was performed (**p<0.01, ***p<0.001, ****p<0.0001).

**Figure 4.**
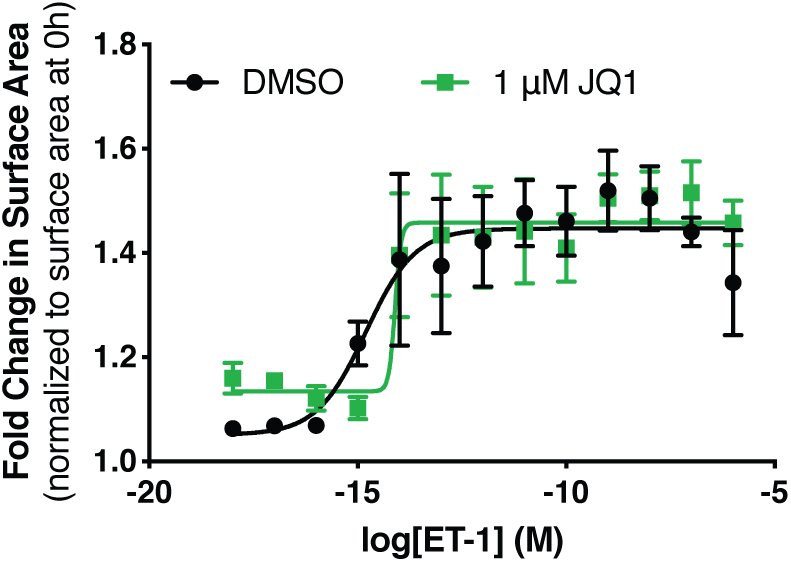
ETR-mediated hypertrophy is insensitive to BET inhibition independent of ET-1 concentration. Dose response curves were generated to assess the effect of JQ1 on cardiomyocyte surface area at a range of ET-1. Cardiomyocytes were treated for 24 h as indicated followed by fixation and staining with Hoechst dye and for α_2_-actinin to identify NCRMs. Fold change in surface area over cardiomyocytes from the same biological replicate at 0 h was determined. Data is presented as mean ± S.E.M for n=3-4 independent experiments. Dose response curves were plotted using sigmoidal dose response (variable slope) curves by non-linear regression.

As JQ1 inhibits all members of the BET family of bromodomain proteins, we confirmed the effects on hypertrophy were mediated by Brd4 and not Brd2 and/or Brd3. Brd2, Brd3, and Brd4 were individually depleted in NRCMs using siRNA (Figure 5A). Hypertrophic responses were then induced through activation of the α_1_-AR or ETR. Brd2 and Brd3 knockdown reduced the basal size of NRCMs relative to control siRNA (Figures 5B), but the response following activation of either receptor was comparable to that observed with control siRNA (Figure 5C). In contrast, Brd4 knockdown recapitulated the effects observed with JQ1 in that it attenuated the response to α_1_-AR activation but did not affect ETR-mediated hypertrophy (Figures 5B and 5C). These data argue that Brd4 inhibition accounts for the receptor-specific effects of JQ1 on cardiomyocyte hypertrophy.

**Figure 5.**
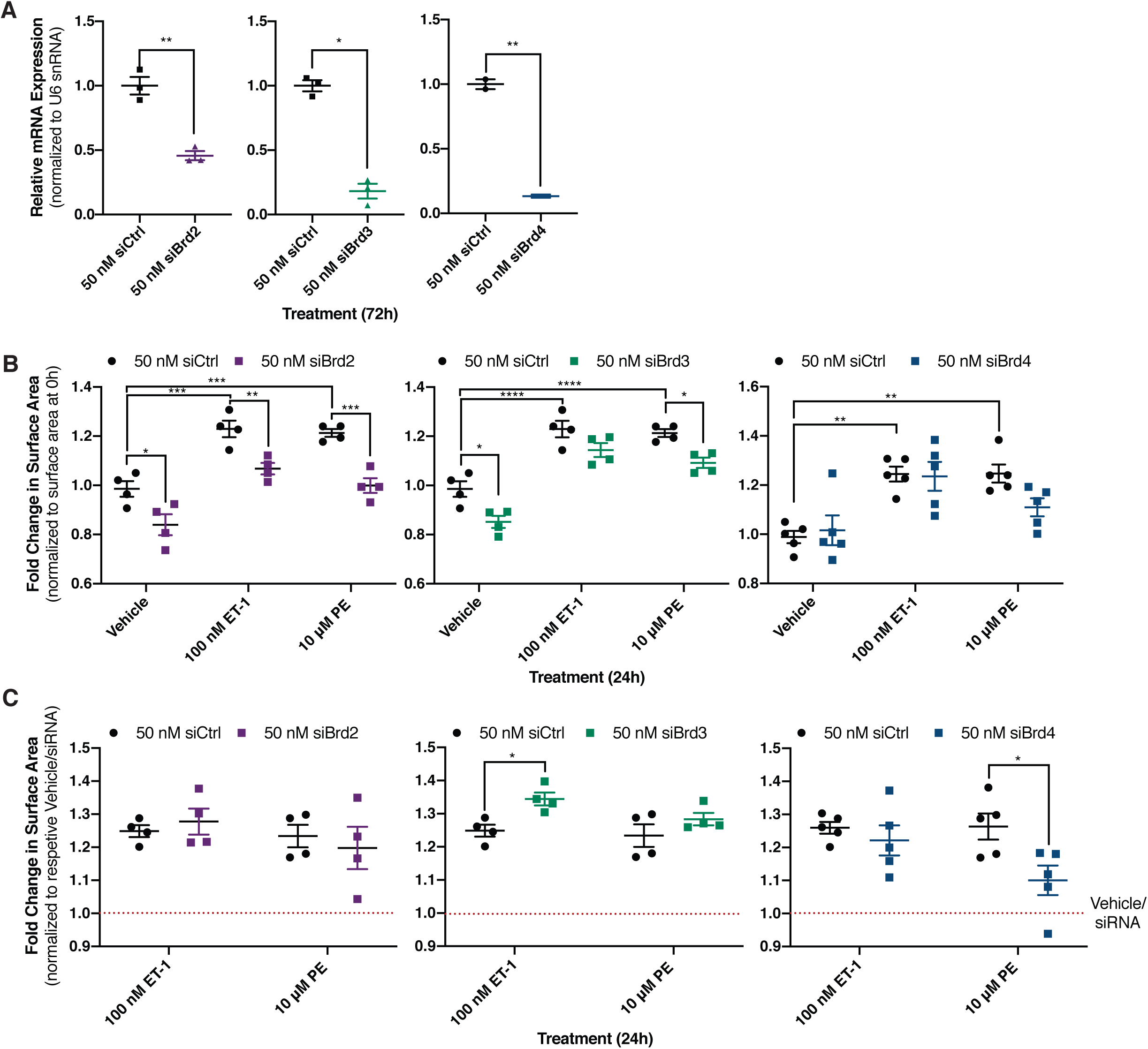
Role of individual BET family members expressed in cardiomyocytes assessed after siRNA-mediated knockdown. **(A)** Brd2, Brd3, and Brd4 knockdown efficiency in NRCMs 72 h after transfection with targeted siRNA was determined by RT-qPCR. An unpaired t-test was performed. **(B)** Fold change in surface area over cardiomyocytes transfected with control siRNA from the same biological replicate at 0 h. Following 72 h knockdown with indicated siRNA, cardiomyocytes were treated for 24 h as indicted. Cardiomyocytes were fixed and identified by staining for the cardiomyocyte specific marker α_2_-actinin. Two-way ANOVA followed by post-hoc t-tests with Bonferroni correction was performed. **(C)** Change in cardiomyocyte size from (B) is presented relative to respective vehicle/siRNA treatment to normalize for the difference in basal size. Data is presented as mean ± S.E.M with each point representing a biological replicate. Two-way ANOVA followed by post-hoc t-tests with Bonferroni correction was performed (*p<0.05, **p<0.01, ***p<0.001).

To gain further insight into how Brd4 function is differentially affected by distinct GPCR signalling pathways, we assessed gene-specific Brd4 localization to chromatin using chromatin immunoprecipitation coupled to qPCR (ChIP-qPCR). ChIP was performed using an antibody recognizing endogenous Brd4 and was quantified by qPCR using primer pairs near the transcription start sites of Nppb, Nppa, and Serpine1. Occupancy at previously defined super enhancers for Nppb and Serpine1 in cardiomyocytes was also assessed (19). At all genomic loci tested, 24 h α_1_-AR activation increased Brd4 chromatin occupancy compared to vehicle treatment (Figure 6A). We observed no change in Brd4 occupancy compared to vehicle following ETR treatment, despite the fact that mRNA levels for the same genes were similarly induced by PE and ET-1 (Figures 2C and 3C). The effects on Brd4 occupancy were not simply a reflection of altered Brd4 protein levels, as immunoblots performed on cell extracts from NRCMs treated with PE or ET-1 revealed slight decreases in expression compared to vehicle controls (Figures 6B and 6C). This suggests that signalling through α_1_-AR, but not ETR, triggers recruitment of Brd4, making gene expression changes downstream of this receptor more sensitive to Brd4 inhibition by JQ1.

**Figure 6.**
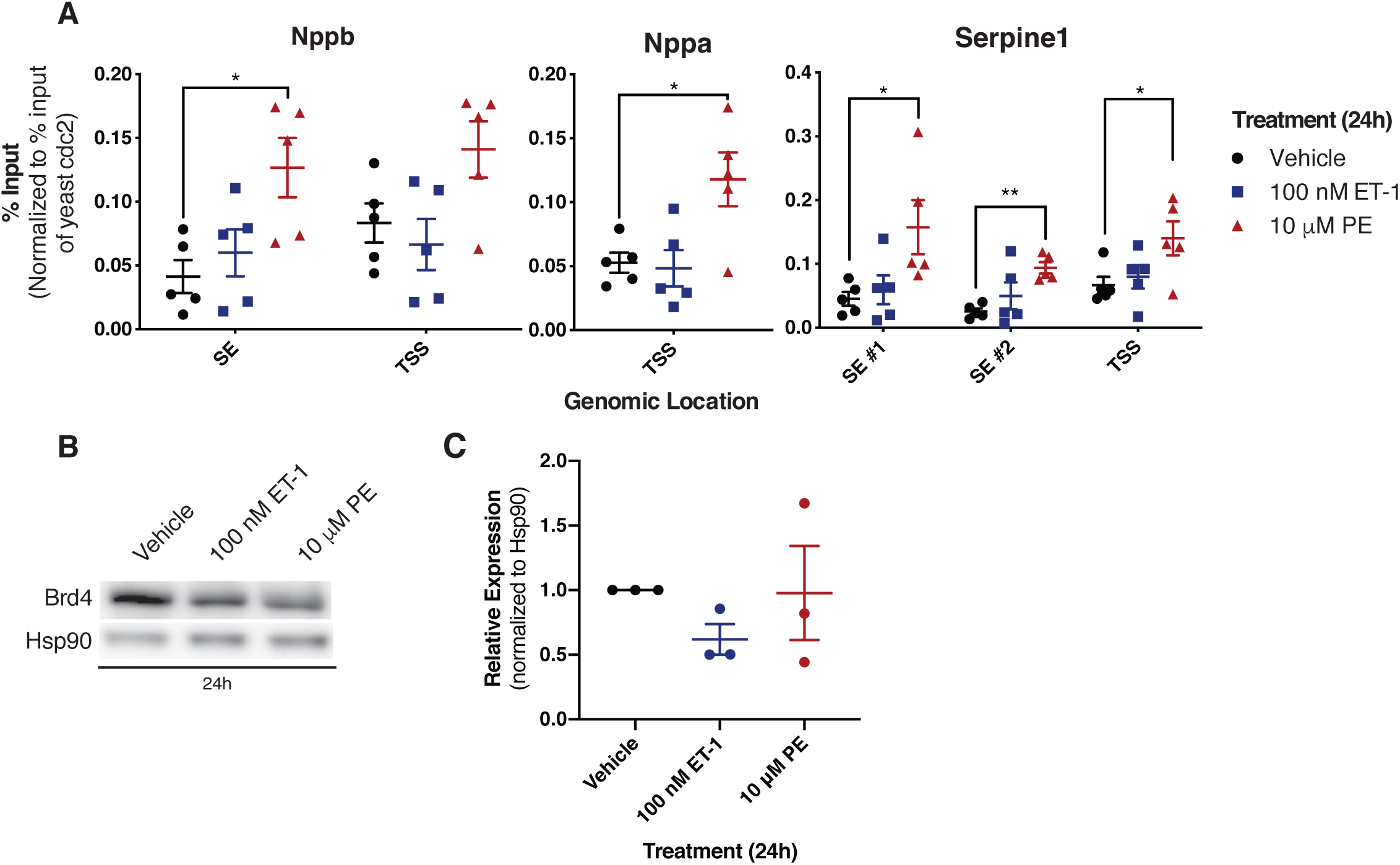
Brd4 chromatin occupancy increases in response to α_1_-AR but not ETR activation. **(A)** Cardiomyocytes were treated for 24 h as indicated. Following treatment, crosslinked chromatin was immunoprecipitated with an anti-Brd4 antibody, followed by DNA purification and quantification by qPCR using primers at the indicated loci. Each immunoprecipitation was normalized to the % input for exogenous *S. pombe* spike-in DNA at the *cdc2^+^* loci. Data was analyzed by one-way ANOVA followed by Dunnett’s post-hoc comparison. **(B)** A western blot to assess changes in Brd4 protein expression following the indicated treatment for 24 h. **(C)** Densitometry based quantification of Brd4 normalized to Hsp90 expression. Data is presented as mean ± S.E.M with each point representing a biological replicate. Data was analyzed by one-way ANOVA followed by Dunnett’s post-hoc comparison (*p<0.05, **p<0.01).

### RNA-seq reveals differences in GPCR-dependent signalling between hypertrophic agonists

To comprehensively profile receptor-specific effects on cardiomyocyte hypertrophy, we performed RNA-seq on NRCMs following activation of either receptor for 24 h in the presence or absence of JQ1. When comparing agonist versus vehicle conditions, we observed robust gene expression changes [log_2_(Fold Change) > ±1 and p-value < 0.05] for hundreds of genes following ETR (209 upregulated, 192 downregulated) or α_1_-AR activation (269 upregulated, 279 downregulated). The genes regulated by either receptor overlapped significantly, although there were more genes uniquely regulated by α_1_-AR activation than by ET-1 activation (Figures 7A and 7B). The combined effect of receptor agonists and JQ1 on differential expression was visualized by performing K-means clustering (Figures 7C and 7D). This analysis identified three major gene clusters that were similarly regulated by agonist and JQ1: one in which genes were repressed by agonist in the presence or absence of JQ1 (cluster 1), one in which genes were activated by agonist and attenuated by JQ1 (cluster 2), and one in which genes were activated by agonist in the presence or absence of JQ1 (cluster 3). The observation that the primary effect of JQ1 was to dampen expression of genes regulated by receptor activation aligns with the known roles of Brd4 in recruiting P-TEFb to regulate pause-release and activate transcription (15), and is consistent with the effect of JQ1 on cardiac stress-induced genes previously characterized (20).

**Figure 7.**
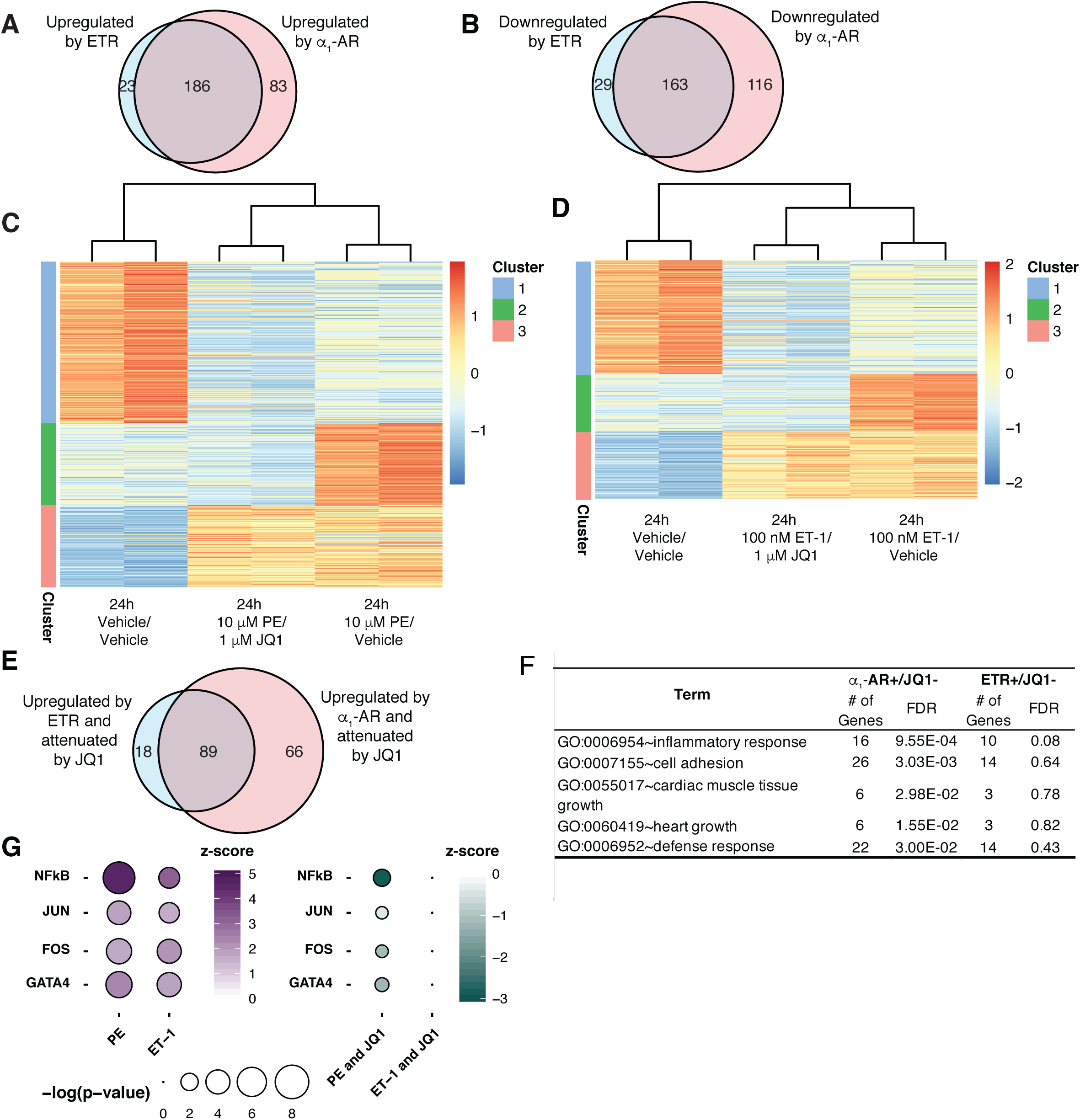
Transcriptome analysis of gene expression programs mediated by receptor activation and the effect of Brd4 inhibition. **(A)** Venn diagram of significantly upregulated genes (log2FC > 1 for Agonist/Vehicle vs Vehicle/Vehicle, p-value < 0.05) or **(B)** downregulated genes (log2FC < −1 for Agonist/Vehicle vs Vehicle/Vehicle, p-value < 0.05) following 24 h receptor activation. **(C and D)** Heat maps were generated for genes differentially regulated following activation of the specified receptor for 24 h. Each row was normalized, with the color representing the z-score for the specific row. K-means clustering was performed to identify subsets of genes with distinct patterns following Brd4 inhibition. **(E)** Venn diagram of genes upregulated by respective receptor activation (log2FC > 1 for Agonist/Vehicle vs Vehicle/Vehicle, p-value < 0.05) and attenuated by Brd4 inhibition from the activated state (log2FC < −0.5 for Agonist/JQ1 vs Agonist/Vehicle, p-value < 0.05). **(F)** Gene ontology enrichment for genes attenuated by JQ1 following receptor activation (from E) performed with DAVID. The false discovery rate (FDR) indicates whether the pathway was significantly enriched in the gene list. **(G)** Changes in transcription factor activity predicted by Ingenuity Pathway Analysis (IPA). The z-score represents the predicted change in transcription factor activity between the two treatment groups and the p-value indicates whether the transcription factor’s targets are significantly enriched in the gene set. The purple dots (left side) indicate the change in activity following agonist treatment alone (log2FC > 1 for Agonist/Vehicle vs Vehicle/Vehicle, p-value < 0.05). The green dots (right side) indicate the JQ1 dependent decrease in activity from the activated state (log2FC < −0.5 for Agonist/JQ1 vs Agonist/Vehicle, p-value < 0.05).

We focused on groups of genes for which increased expression caused by activation of either receptor was attenuated by JQ1 [log_2_(Fold Change) < −0.5 compared to agonist alone; p-value < 0.05]. JQ1 attenuated expression of 107 ETR-induced genes and 155 α_1_-AR-induced genes (termed receptor+/JQ1-) (Figure 7E). A majority of the ETR+/JQ1-genes overlapped with α_1_-AR+/JQ1-genes (83.2% overlap, 16.8% unique), whereas there was a large number of unique α_1_-AR+/JQ1-genes (57.4% overlap, 42.6% unique). Gene ontology term analysis of these gene sets revealed that the terms inflammatory response, defense response, cell adhesion, cardiac muscle tissue growth and heart growth, which correspond to the pathophysiology of cardiomyocyte hypertrophy and align with those previously identified to be affected by JQ1, were significantly enriched among the AR+/JQ1-genes (Figure 7F) (20). In contrast, none of these terms were significantly enriched among ETR+/JQ1-genes, consistent with the selective effect of JQ1 on the α_1_-AR response.

Previous studies have identified multiple transcription factors that are required to activate pro-hypertrophic genes in cardiomyocytes (9, 34). Some of these transcription factors have been associated with Brd4 activity in cardiomyocytes, either through motif enrichment in genomic loci with high Brd4 occupancy or various gene set enrichment methods for JQ1-sensitive genes. The pathway-specific effect of Brd4 inhibition suggests that specific transcription factors may be dependent on Brd4 to activate transcription. We initially focused on those transcription factors that were previously implicated including, NF-κB, GATA4, and the AP-1 subunits Jun and Fos (Figure 7G) (18–20). To determine the effect of JQ1 on these transcription factors, we used Ingenuity Pathway Analysis software (IPA; Qiagen) to predict changes in their activity (Figure 7G). IPA upstream regulator analysis provides a z-score, to indicate the predicted change in activity between the two treatment groups, and a Fisher’s exact test p-value, to indicate whether particular upstream regulator’s target genes are significantly enriched in the gene expression program. Interestingly, IPA predicted increased activity of these transcription factors following activation of either receptor (positive z-score, agonist/vehicle vs vehicle/vehicle), but activity was specifically attenuated by JQ1 following α_1_-AR activation (negative z-score, agonist/JQ1 vs agonist/vehicle) (Figure 7G). This suggests that receptor-specific activation mechanisms for these transcription factors dictate their dependence on Brd4 activity. When we expanded the analysis to include all activated transcription factors a more ubiquitous effect of Brd4 inhibition was observed. We identified 78 transcription factors with enhanced activity following α_1_-AR activation of which activity of 39 (50%) were attenuated by co-treatment with JQ1. In contrast, ETR activation was predicted to enhance activity of 50 transcription factors and only eight (16%) were attenuated by JQ1. Thus, although specific transcription factors may function to recruit Brd4 to specific loci, the more ubiquitous effect of Brd4 inhibition on α_1_-AR-mediated transcription factor activity suggests that α_1_-AR signalling may increase the pool of active Brd4.

### Signalling pathway regulating Brd4 recruitment to chromatin involves PKA

We hypothesized that a distinct signalling pathway activated by the α_1_-AR determines differential recruitment of Brd4 and inhibitory effect of JQ1 on transcription factor activity. Brd4 is activated following phosphorylation of its phosphorylation dependent interaction domain (PDID). An *in vitro* kinase assay demonstrated that PKA was able to phosphorylate this region, although the functional significance was not determined (35). We have previously demonstrated that α_1_-AR, but not ETR activation, led to Gαs-dependent activation of cAMP/PKA signalling in HEK 293 cells (36). We thus hypothesized that PKA, a protein kinase activated by cAMP, regulates the specific effects of α_1_-AR signalling on Brd4 in cardiomyocytes.

We confirmed that α_1_-AR and not ETR signalling activated PKA in NRCMs. Cardiomyocytes were transduced with a nuclear-localized Förster resonance energy transfer (FRET)-based PKA biosensor (AKAR4-NLS) to monitor PKA activity (37). When phosphorylated, the biosensor undergoes a conformational change that moves the two fluorophores into closer proximity, leading to an increased FRET ratio. Nuclear localization of the AKAR biosensor was confirmed by fluorescence microscopy (Figure 8A). We then generated a dose-response relationship for PKA activity following stimulation with ET-1 and PE. Alprenolol was included in the PE experiments to prevent off-target effects on the β-AR at high concentrations. We observed a dose-dependent increase in FRET, indicating an increase in PKA activity, following 15 min α_1_-AR activation. In contrast, no change in activity was observed following 15 min ETR activation (Figure 8B). This demonstrated that the α_1_-AR uniquely activates PKA in the nucleus of cardiomyocytes, similar to what we detected in HEK 293 cells (36).

**Figure 8.**
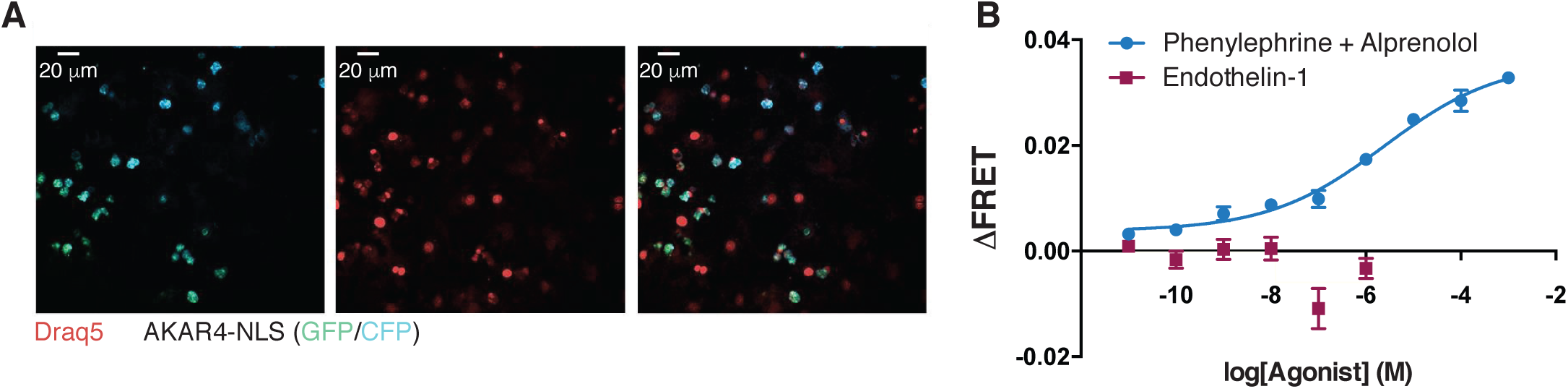
α_1_-AR activation leads to increased nuclear PKA signalling. **(A)** Cardiomyocytes were transduced with AAV9-AKAR4-NLS virus at a MOI of 5000 and imaged 72 h later. Nuclei were visualized by staining live cells with Draq5. **(B)** Dose-response curves for PKA activation following activation of the ETR or α_1_-AR were generated. Alprenolol was included to prevent off-target β-AR activation by high concentrations of PE. Data is presented mean ± S.E.M for three biological replicates. Dose response curves were plotted using sigmoidal dose response (variable slope) curves by non-linear regression.

In order to determine if PKA activity downstream of the α_1_-AR regulates Brd4 function in cardiomyocytes, we performed ChIP-qPCR after inhibition or activation of PKA. To inhibit PKA, we used the competitive inhibitor KT5720 (38). As we anticipated that long-term PKA inhibition could cause other changes in cellular physiology that would complicate interpretation of the experiments, we used a treatment time of 1.5 h. We quantified Brd4 localization using primer pairs near the transcription start site of c-Fos and Ctgf as well as previously defined super-enhancer regions of Ctgf (19). Activation of the α_1_-AR for 1.5 h enhanced Brd4 occupancy near the transcription start sites of both genes and along the previously defined Ctgf super enhancers, whereas ETR stimulation had no effect (Figures 9A and 9B). These data suggest that the specific effect of α_1_-AR signalling on Brd4 chromatin occupancy is maintained at the shorter treatment time. Pre-treatment with KT5720 abrogated the increase in Brd4 occupancy following α_1_-AR activation, consistent with a requirement for PKA activity for chromatin recruitment of Brd4 downstream of α_1_-AR signalling (Figure 9A and 9B). PKA inhibition also increased the basal occupancy of Brd4 at these sites, perhaps reflecting a repressive function for PKA in unstimulated NRCMs. We also stimulated activation of PKA in NRCMs by increasing intracellular cAMP levels with forskolin and IBMX, an adenylyl cyclase activator and phosphodiesterase inhibitor, respectively (39). Sustained PKA activation (1.5 h) increased Brd4 chromatin association at two of the four genomic loci assessed, reinforcing the key role of PKA in Brd4 activation (Figure 9C).

**Figure 9.**
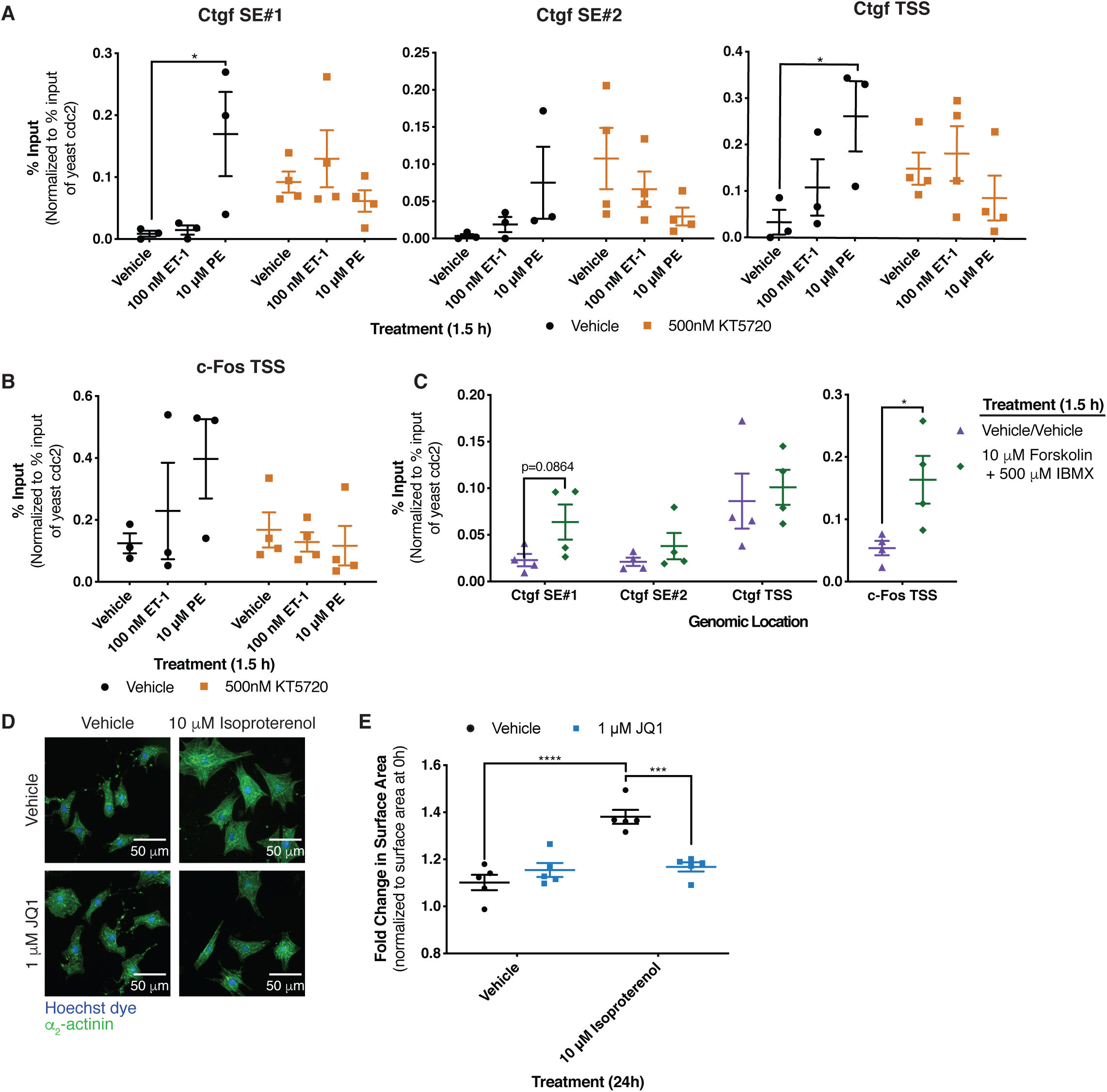
PKA signalling regulates recruitment of Brd4 to chromatin. **(A and B)** Effect of PKA inhibition with the small-molecule inhibitor KT5720 on receptor-mediated increases in Brd4 occupancy. Cardiomyocytes were pre-treated for 30 min with the PKA inhibitor prior to receptor activation for 1.5 h with the indicated agonists. Two-way ANOVA followed by post-hoc t-test comparisons with Bonferroni correction was performed. **(C)** PKA was activated for 1.5 h by increasing intracellular cAMP levels with forskolin and IBMX, an adenylyl cyclase activator and phosphodiesterase inhibitor respectively. Following indicated treatment, cardiomyocytes were fixed, and ChIP was performed with an anti-Brd4 antibody. ChIP was quantified by qPCR using primers at the indicated loci. An unpaired t-test was performed. **(D)** After 24 h of the indicated treatments, NRCMs were fixed and stained with Hoechst dye and for the cardiomyocyte specific marker α_2_-actinin. **(E)** Fold change in surface area after 24 h of the indicated treatment over cardiomyocytes fixed at 0 h from the same biological replicate. Two-way ANOVA followed by post-hoc t-test comparisons with Bonferroni correction was performed. Data is presented as mean ± S.E.M with each point representing a separate biological replicate. (*p<0.05, ***p<0.001, ****p<0.0001).

To test whether a regulatory link between PKA and Brd4 could be detected in response to other GPCRs coupled to Gαs, we examined the role of Brd4 downstream of the β-AR, activation of which is strongly pro-hypertrophic in cardiomyocytes (40). The primary signalling pathway downstream of this receptor in cardiomyocytes (and other cell types as well) involves adenylyl cyclase activation, cAMP production, and increased protein kinase A activity (41). We thus predicted that hypertrophy mediated by the β-AR would also be attenuated by inhibition of Brd4 with JQ1. Following 24 h treatment with the agonist isoproterenol, we observed a ∼25% increase in surface area which was completely blocked by co-treatment with JQ1 (Figure 9D and 9E). This demonstrates that the observed connection between PKA and Brd4 is not unique to α_1_-AR signalling and may reflect a general Gαs-coupled GPCR-dependent pathway for Brd4 activation. Taken together, our results point to PKA and Brd4 as central players underlying receptor-specific gene regulatory mechanisms in hypertrophic cardiomyocytes (Figure 10).

**Figure 10.**
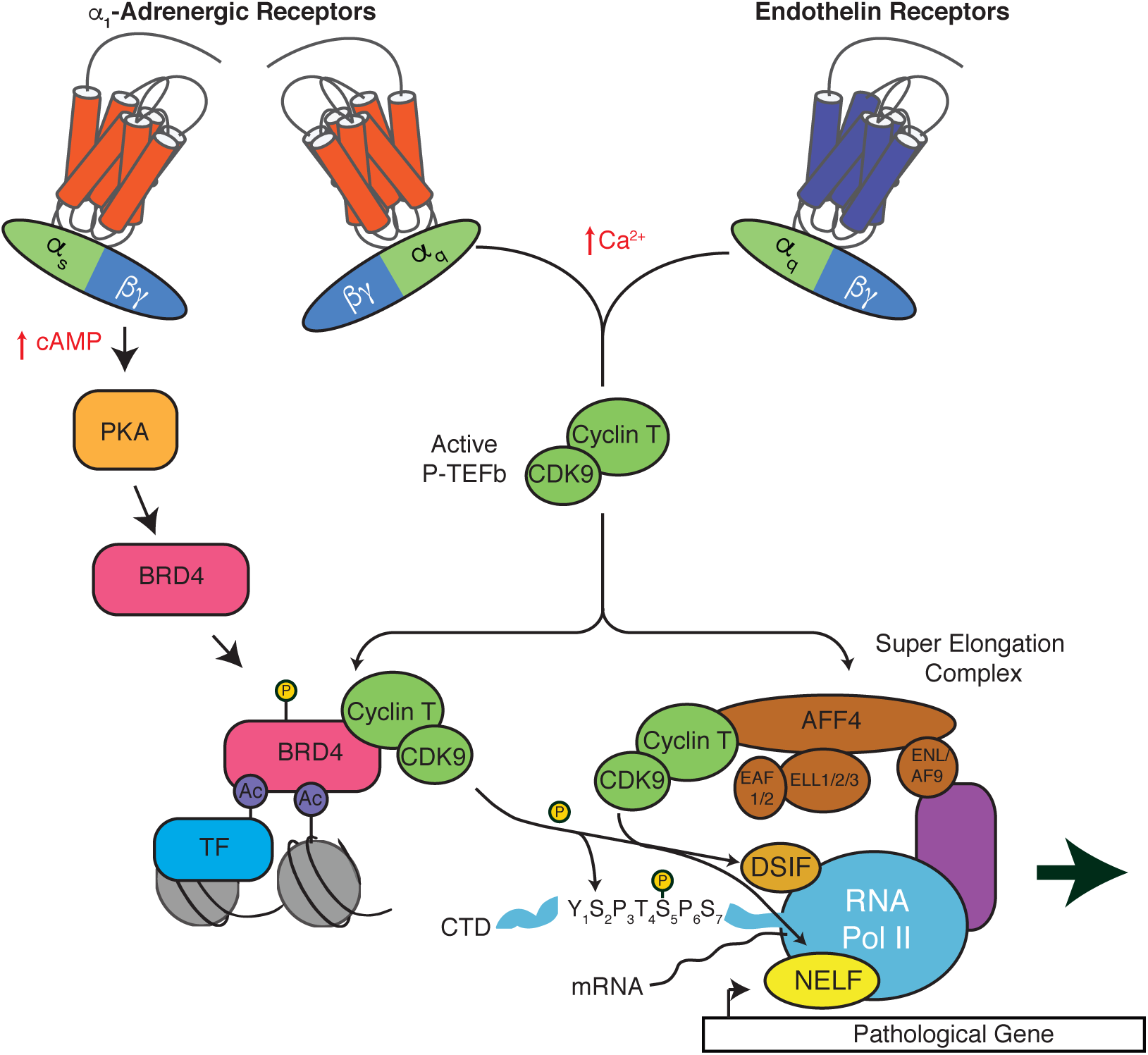
Model of P-TEFb complex activation following activation of the α_1_-AR or ETR. Both the ETR and α_1_-AR activate a signalling cascade which increases active P-TEFb and requires subsequent recruitment through the SEC. The α_1_-AR activation also leads to Brd4-dependent recruitment due to the activation of a PKA signalling pathway.

## Discussion

In this study, we showed that activation of distinct GPCRs in cardiomyocytes can result in hypertrophic responses and gene expression programs that are qualitatively similar, but that operate through different transcriptional regulatory mechanisms. We also identified a novel mechanism regulating Brd4 in the development of cardiomyocyte hypertrophy that expands our understanding of how specific signalling pathways regulate the recruitment of general transcription regulators and that may have relevance in other physiological contexts.

Chromatin occupancy of Brd4 undergoes extensive redistribution in order to positively regulate expression of the cardiomyocyte hypertrophic gene program. These changes lead to enhanced Brd4 occupancy on specific super-enhancers and promoter regions. Previous Brd4 ChIP-seq experiments have used primary cardiomyocytes treated with PE (in accord with our results), or cardiac tissue isolated from mice subjected to transverse aortic constriction (TAC) (19, 20). The increased genomic loading of Brd4 in heart failure models has been attributed to increases in Brd4 expression (19), similar to proposed mechanisms in various forms of cancer (42–44). In heart failure, increased Brd4 protein expression is thought to occur due to decreased expression of the Brd4 targeting microRNA miR-9 (21, 45). These reports assessed whole cardiac tissue not enriched for cardiomyocytes or *in vitro* cardiomyocyte studies which reported conflicting evidence regarding changes in Brd4 expression (18, 21). Following 24 h activation of the α1-AR or ETR in cardiomyocytes, we did not observe a significant change in Brd4 protein expression (Figure 6B). Instead, the increased Brd4 recruitment following α_1_-AR activation was dependent on cAMP/PKA signalling pathway.

Phosphorylation of Brd4 is critical for its activation and also mediates alterations in its interactome. Brd4 hyperphosphorylation correlates with its oncogenic potential and increased phosphorylation levels lead to development of BET inhibitor resistance in certain types of cancer (46, 47). At present, casein kinase 2 (CK2) is the only protein kinase demonstrated to directly phosphorylate Brd4, although others have been shown to regulate Brd4 activity (35, 46, 48). CK2 phosphorylation sites reside within the PDID domain, where phosphorylation results in a conformational change that unmasks the second bromodomain and enables interactions with acetyl-lysine residues (35). An *in vitro* kinase screen of the PDID domain identified PKA as a potential Brd4 kinase, aligning with the PKA regulatory effect on Brd4 we observed (35). This suggests that in cardiomyocytes, enhanced PKA activity could increase PDID phosphorylation and drive the conformational change required for Brd4 to interact with chromatin. Further work remains to identify the putative phosphorylated sites and elucidate their functional role(s). Conversely, we also observed an increase in basal Brd4 occupancy following PKA inhibition (Figure 9A). Such increased Brd4 occupancy may be related to PKA’s known role in regulating histone deacetylases (49). The balance between these two opposing processes regulated by PKA is likely linked to the highly localized nature of PKA signalling through interactions with A-kinase anchoring proteins (AKAPs) (50).

RNA-seq analyses revealed that the transcriptional programs triggered by activation of α_1_-AR or ETR were highly overlapping, consistent with the similar hypertrophic response downstream of either receptor. The mechanistic difference between the two pathways was instead linked to the fact that specific groups of genes relevant to the hypertrophic response were differentially sensitive to JQ1 downstream of α_1_-AR activation compared to ETR. Previous reports have identified inflammatory pathways enriched in JQ1-attenuated genes using both *in vitro* and *in vivo* models of cardiomyocyte hypertrophy (20). We observed JQ1-selective attenuation of these inflammatory pathways following α_1_-AR activation (Figure 7F). We argue that this difference stems from a greater role for Brd4 downstream of α_1_-AR, consistent with our Brd4 ChIP results. Such inflammatory responses are characteristic of heart failure, and the production of several cytokines is increased during cardiac remodelling (51, 52). Importantly, inflammation is a driver of cardiomyocyte hypertrophy and inhibition of those pathways prevents the progression of heart failure (53, 54). Although specific pathways were not enriched among genes attenuated by JQ1 follow ETR activation, a large number of genes did exhibit JQ1 sensitivity. We suspect the observed effects are likely due to functions of Brd2 and/or Brd3 in regulating these genes, although further work is required to confirm this.

Further evidence for the effect of JQ1 on the inflammatory response is evident by the negative effect on inflammatory transcription factors such as NF-κB and AP-1 (Figure 7G) (34, 55, 56). We focused on these transcription factors for two reasons: their gene expression signatures were previously identified in JQ1-sensitive, TAC-induced genes, or a causal role in regulating Brd4 recruitment in cardiomyocytes has been identified (18, 19). Importantly, these transcription factors are also directly implicated in driving pathological cardiomyocyte hypertrophy in various *in vitro* and *in vivo* models (57–61). Despite the fact that similar transcription factors were activated downstream of α_1_-AR and ETR, JQ1 only attenuated the α_1_-AR response. The receptor-specific attenuation of transcription factor activity may be due to distinct signalling mechanisms required for transcription factor activation, creating a differential dependence on Brd4 activity (62–65). However, the greater number of transcription factors attenuated by JQ1 following α_1_-AR activation suggests that it leads to an active form of Brd4 that more readily binds to chromatin and promotes the activity of transcription factors. Although we predict that PKA activates Brd4 directly, indirect activation through a downstream factor is also possible. Further investigation is required to determine the effects of JQ1 on Brd4’s interactome and phosphorylation status following α_1_-AR or ETR activation.

Receptor-specific activation of Brd4 we identified may have implications for the clinical use of Brd4 inhibitors in cardiovascular disease. Efficacy will be dependent on the specific neurohormonal signalling pathways altered in a particular patient. Specifically, we would expect a negative correlation with a patient’s ET-1 levels. Therefore, more extensive characterization of signalling molecules in patients might be important predictors of drug efficacy. Importantly, we expect that Brd4 inhibition will be less effective in severe and/or late stages of heart failure as PKA activity is reduced (41) and levels of endothelin-1 or its precursors (66–68) increases. Chronic infusion of these neurohormones with mice is required to determine how the receptor-specific effects of Brd4 inhibition effects cardiac remodelling. Furthermore, although JQ1 has been demonstrated to reverse established heart failure in mouse TAC and myocardial infarction mouse models (20), we expect JQ1 efficacy would decrease as heart failure progresses.

Our finding that JQ1 sensitivity is dependent on activation of PKA signalling raises the question of whether Brd4 inhibition is also an effective therapeutic for other pathologies associated with enhanced PKA signalling. For example, chronic activation of PKA is a hallmark of dopamine-dependent neuronal pathologies such as cocaine addiction and L-DOPA induced dyskinesia (LID) (69, 70). Recent studies have implicated Brd4 in regulating the neuronal transcriptional programs and behavioural effects driven by dopamine signalling in these contexts. Systemic administration of JQ1 reduces reward seeking behaviour in addiction models and prevents LID development in Parkinson’s models (71, 72). The correlation between PKA and Brd4 activity in these cases suggests that the regulation of Brd4 chromatin occupancy by PKA might be a common regulatory mechanism for other GPCRs and cell types. Furthermore, certain adrenocortical adenomas are driven by enhanced basal PKA activity due to activating mutations in Gαs or the catalytic subunit of PKA (73, 74). We expect these adenomas would be highly sensitive to Brd4 inhibition, although further work is required to establish the requirement of Brd4 in progression of these cancers.

Dysregulated activation and recruitment of P-TEFb is an underlying cause of several diseases and developmental disorders (75). While recruitment of P-TEFb is regulated either by Brd4 or the SEC, little is known about the functional relationship between these two complexes. It has been suggested that these complexes work together to target P-TEFb to different substrates whereas others have shown that Brd4 assists in recruiting the SEC (76, 77). Our results indicate the cooperative nature of these two complexes is dependent on the signalling pathway employed to activate transcriptional responses. Following ETR activation, the SEC alone is sufficient to elicit a gene expression program required for cardiomyocyte hypertrophy, whereas the activation of the Gαs/cAMP/PKA pathway by the α_1-_AR leads to an additional dependence on Brd4. This suggests that the SEC form of P-TEFb may have a general transcriptional regulatory role whereas the form associated with Brd4 may have a more restricted signal-responsive role. Further investigation is required to assess whether these complexes have distinct or overlapping functions in cardiomyocytes. Furthermore, expanding our understanding of how signalling pathways activate these complexes is an important step to improve therapeutic approaches in diseases with dysregulated P-TEFb.

## Methods

### Primary neonatal rat cardiomyocyte isolation, tissue culture, transfection and treatments

Unless otherwise stated, all reagents were obtained from Sigma. Primary rat cardiomyocytes were isolated from 1-3 day old Sprague-Dawley rats (Charles River Laboratories, St-Constant, Quebec) as previously described with minor modifications (78). Following isolation, cardiomyocytes were seeded at a density of 40 000 cells/cm^2^ on tissue culture dishes coated with 0.1% gelatin and 10 μg/mL fibronectin in DMEM low glucose (Wisent) supplemented with 7% FBS (Wisent) (v/v), 1% (v/v) penicillin/streptomycin (P/S), and 10 μM cytosine-β-d-arabinoside (MP Biomedicals). After 24 h, plates were washed twice with DMEM low glucose and media changed to cardiomyocyte maintenance media (DMEM low glucose, 1% (v/v) insulin/selenium/transferrin (Wisent), and 1% (v/v) P/S) with 10 μM cytosine-β-d-arabinoside. Twenty-four hours later, media was replaced with fresh cardiomyocyte maintenance media and experiments were initiated 24h later. Cardiomyocytes were maintained at 37°C with 5% CO_2_ and typical cultures contained >90% cardiomyocytes. Cardiomyocytes were treated with endothelin-1 (Bachem), phenylephrine, iCdk9 (Novartis), KL-2 (ProbeChem Biochemicals), JQ1, alprenolol, KT5720, forskolin, 3-isobutyl-1-methylxanthine (IBMX), or isoproterenol.

For small interfering RNA (siRNA) transfection (siGENOME SMARTPool, Horizon Discover), cardiomyocytes were pelleted at 400 g for 5 min at 4°C after isolation, resuspended in DMEM low glucose supplemented with 2.5% (v/v) FBS and plated at a density of 60 000 cells/cm^2^. Cardiomyocytes were transfected with 50 nM siRNA for the specified target gene with Lipofectamine 2000 (Invitrogen) according to the manufacturer’s instructions. After 5h incubation, the media was replaced with DMEM low glucose supplemented with 7% FBS (v/v), 1% (v/v) penicillin/streptomycin (P/S), and 10 μM cytosine-β-d-arabinoside and cultured as previously described.

### Immunofluorescence and measurement of cell area

Cardiomyocytes were plated in 96-well plates and cultured as described. Following indicated treatment, cells were fixed with methanol for 5 min at −20°C, permeabilized with 0.2% Triton X-100 (v/v) in PBS for 5 min at room temperature and blocked with 10% horse serum (Wisent) in PBS for 1 h at room temperature. Primary anti-α_2_-actinin antibody (Sigma, A7811; 1/200) in 10% horse serum/PBS was incubated with cardiomyocytes overnight at 4°C. The following day, the fixed cardiomyocytes were incubated with anti-mouse Alexa Fluor 488 secondary antibody (Invitrogen, A-11029; 1/500) in 10% horse serum/PBS for 1 h at room temperature and 10 min with Hoechst dye (Invitrogen) (1 μg/μL) in PBS at room temperature. Stained cardiomyocytes were imaged with an Operetta high-content screening system (PerkinElmer) with 20X magnification and analyzed with Columbus Image Analysis System (PerkinElmer). Hoechst dye was excited with a 360-400 nm filter and emissions detected at 410-480 nm and Alexa 488 was excited with a 460-490 nm filter and emissions detected at 500-550 nm. The average of two technical replicates was taken for all treatments.

### AKAR4-NLS AAV transduction and FRET experiments

The AKAR4-NLS construct was a gift from Dr. Jin Zhang (UCSD) and the pAAV-CAG-GFP was a gift from Dr. Karel Svoboda (Addgene plasmid #28014) (79, 80). To generate the AKAR4-NLS biosensor for adeno-associated virus (AAV) production, AKAR4-NLS biosensor was excised and cloned into the pAAV backbone using BamHI and EcoRI (New England Biolabs) at the 5’ and 3’ end, respectively. For AAV production, HEK 293T cells were maintained in DMEM high glucose supplemented with 10% (v/v) FBS and 1% (v/v) P/S in a controlled environment of 37°C and 5% CO_2_. Adeno-associated viruses were produced as previously described (81).

Twenty-four hours after plating cardiomyocytes, media was changed to cardiomyocyte maintenance media with AAV9-packaged AKAR4-NLS biosensor at a multiplicity of infection (MOI) of 5000. Following 24 h transduction, media was changed to cardiomyocyte maintenance media and changed every 24 h until experiment. After 48 h incubation, cardiomyocyte maintenance media was removed and cells were washed with Krebs solution (146 mM NaCl, 4.2 mM KCl, 0.5 mM MgCl_2_, 1 mM CaCl_2_, 10 mM HEPES pH 7.4, 1 g/L glucose) and incubated for 1 h at 37°C with 5% CO_2_ in Krebs solution prior to FRET readings. All cardiomyocyte FRET experiments were performed using the Opera Phenix^™^ High Content Screening System (PerkinElmer) with the confocal setting at 40X magnification at 37°C and 5% CO_2_ and analyzed with Columbus Image Analysis System (PerkinElmer). Each well was excited with 425 nm light and emissions detected at 434-515 nm for CFP and 500-550 nm for YFP. Basal FRET images were obtained prior to addition of agonist and stimulated FRET images were obtained 15 minutes after addition of agonist to indicated final concentration. For experiments requiring a β-AR antagonist, 1 μM alprenolol was added to cardiomyocytes 30 minutes prior to obtaining basal FRET images. The FRET ratio was calculated as YFP emission/CFP emission. For all experiments, ΔFRET refers to: (Stimulated Agonist FRET Ratio – Basal Agonist FRET Ratio) – (Stimulated vehicle FRET Ratio – Basal FRET Ratio). The average of three technical replicates was taken for all treatments.

Following FRET experiments, cardiomyocytes were stained with 5 μM Draq5 at room temperature for 5 minutes. Images were obtained on the Opera Phenix^™^ High Content Screening System (PerkinElmer) using the confocal setting at 40X magnification. Draq5 was imaged using 640 nm excitation and emissions detected at 650-760 nm, YFP using 425 nm excitation and emission detected at 434-515 nm, and CFP using 425 nm excitation and emission detected at 500-550 nm.

### RT-qPCR

Following indicated treatments of cardiomyocytes, cells were lysed in TRI reagent and RNA was extracted following the manufacturer’s protocol. Reverse transcription was performed with random hexamer primers using an MMLV-RT platform (Promega) according to the manufacturer’s protocol. Subsequent qPCR analysis was performed with BrightGreen 2x qPCR Master mix (Applied Biological Materials Inc.) on a Bio-Rad 1000 Series Thermal Cycling CFX96 Optical Reaction module. Ct values were normalized to U6 snRNA and fold change over respective control was calculated using 2^-**ΔΔ**Ct^ method. Primer sequences were the following: Nppb (5’ CAATCCACGATGCAGAAGCTG 3’ and 5’ TTTTGTAGGGCCTTGGTCCTTT 3’), Nppa (5’ CCTGGACTGGGGAAGTCAAC 3’ and 5’ ATCTATCGGAGGGGTCCCAG 3’), Serpine1 (5’ TCCTCGGTGCTGGCTATGCT 3’ and 5’ TGGAGAGCTTTCGGAGGGCA 3’), and U6 snRNA (5’ TGGAACGATACAGAGAAGATTAG 3’ and 5’ GAATTTGCGTGTCATCCTTG 3’).

### RNA-seq analysis

RNA was isolated with the RNeasy® Mini Kit (Qiagen) according to manufacturer’s instructions. Libraries were prepared using the NEBNext^®^ rRNA-depleted (HMR) stranded library kit and single-read 50bp sequencing completed on the Illumina HiSeq 4000 at the McGill University and Génome Québec Innovation Centre, Montréal, Canada. Reads were trimmed with TrimGalore (0.6.0) (82, 83) using the following settings: --phred33 --length 36 -q 5 –stringency 1 -e 0.1. Following processing, reads were aligned to the Ensembl rat reference genome (Rattus_norvegicus.Rnor_6.0.94) (84) with STAR (2.7.1a) (85). Transcripts were assembled with StringTie (1.3.4d) (86) and imported into R (3.6.1) with tximport (1.12.3) (87). Differential gene expression was assessed with DESeq2 (1.24.0) (88) with the independent hypothesis weighting (IHW) library for multiple testing adjustment (89). Heatmaps and K-means clustering was completed with pheatmap and the removeBatchEffect function from limma (3.40.6) (90) was used prior to data visualization. Pathway analysis was completed with Ingenuity Pathway Analysis *(IPA, QIAGEN Inc.,* https://www.qiagenbio-informatics.com/products/ingenuity-pathway-analysis) (91).

### Chromatin immunoprecipitation-qPCR

Preparation and immunoprecipitation of cardiomyocyte chromatin was performed as previously described, with minor modifications (92). Following the indicated treatments, cardiomyocytes were crosslinked with 1% formaldehyde in DMEM low glucose for 10 min at room temperature with slight agitation. Crosslinking was quenched by addition of glycine to 125 mM final concentration and incubated for 5 min at room temperature with slight agitation. Cardiomyocytes were placed on ice following fixation, washed once with cold PBS, scraped into PBS with 1 mM PMSF and pelleted at 800 g for 5 min at 4°C. The pellet was resuspended in lysis buffer (10 mM Tris-HCl pH 8.0, 10 mM EDTA, 0.5 mM EGTA, 0.25% Triton X-100, 1 mM PMSF, 1x protease inhibitor cocktail) and incubated for 10 min at 4°C on a nutator. Nuclei were pelleted at 800 g for 5 min at 4°C and resuspended in nuclei lysis buffer (50 mM TrisHCl pH 8.0, 10 mM EDTA, 1% SDS, 1 mM PMSF, 1x protease inhibitor cocktail). Nuclei were incubated for 15 min on ice followed by sonication with a BioRuptor (Diagenode) (18 cycles, 30 s on/off, high power). Insoluble cellular debris was removed by centrifugation at 14 000 g for 10 min at 4°C. A small aliquot was taken for quantification and the remaining sample stored at −80°C until use. The aliquot was incubated at 65°C overnight to reverse crosslinks, treated with RNase A (50 μg/mL) for 15 min at 37°C, and then treated with proteinase K (200 μg/mL) for 1.5 h at 42°C. Protein was removed by phenol/chloroform extraction and DNA precipitated at −80°C with 0.3M sodium acetate pH 5.2, 2.5 volumes of 100% ethanol, and 20 µg of glycogen. Samples were centrifuged for 20 min at 16 000 g, the pellet was washed with 70% ethanol, resuspended with ddH_2_O and quantified using a NanoDrop spectrophotometer (Thermo Fisher) to determine concentration of chromatin for each sample.

For immunoprecipitations, 10 μg of chromatin was diluted 9x with dilution buffer (16.7 mM Tris-HCl pH 8.0, 1.2 mM EDTA, 167 mM NaCl, 0.01% SDS, 1.1% Triton X-100, 1x protease inhibitor cocktail). *S. pombe* chromatin, prepared as previously described (93), was spiked-in to each sample for normalization. A rabbit anti-Brd4 antibody (Bethyl, A301-985A; 5ug) or rabbit IgG antibody (Millipore, 12-370; 5 μg), as well as anti-*S. pombe* H2B antibody (Abcam, ab188271), were added to respective IPs and 1% input sample taken for subsequent analysis. Each IP was incubated at 4°C overnight on a nutator, followed by addition of 15 μL Protein G Dynabeads (Invitrogen) in dilution buffer for 4 h. Beads were washed 2X with low salt buffer (20 mM Tris-HCl pH 8.0, 2 mM EDTA, 150 mM NaCl, 0.1% SDS, 1% Triton X-100), 2X with high salt buffer (20 mM Tris-HCl pH 8.0, 2 mM EDTA, 500 mM NaCl, 0.1% SDS, 1% Triton X-100), 1X with LiCl buffer (10 mM Tris pH 8.0, 1 mM EDTA, 0.25M LiCl, 1% NP-40, 1% deoxycholate), 1X with TE buffer (10mM Tris-HCl pH 8.0, 1 mM EDTA) at 4°C. Beads were resuspended in elution buffer (200 mM NaCl, 1% (w/v) SDS) and heated at 65°C for 20 min to elute chromatin. The eluted chromatin was incubated at 65°C overnight to reverse crosslinks and then incubated with proteinase K (200 μg/mL) for 2 h at 37°C. DNA was purified and quantified as described above.

Localization was assessed by qPCR with primers for specific genomic loci; a primer pair amplifying *S. pombe cdc2^+^* was used for normalization. All qPCR reactions were performed using a Bio-Rad 1000 Series Thermal Cycling CFX96 Optical Reaction module and iQ SYBR Green Supermix (Bio-Rad). For each primer pair in a given experimental condition, percent input for IgG control IP was subtracted from the percent input for the Brd4 IP, followed by normalization to the percent input of *S. pombe cdc2^+^*. Primer sequences were the following: Nppb SE (chr5:164778453-164778528, 5’ AGGTGGCACCCCCTCTTCTAC 3’ and 5’ TTGGGGGAGTCTCAGCAGCTT 3’), Nppb TSS (chr5:164796330-164796402, 5’ TTTCCTTAATCTGTCGCCGC 3’ and 5’ GGATTGTTCTGGAGACTGGC 3’), Nppa TSS (chr5:164808403-164808456, 5’ GTGACGGACAAAGGCTGAGA 3’ and 5’ ATGTTTGCTGTCTCGGCTCA 3’), Serpine1 SE #1 (chr12:22636488-22636538, 5’ TCCCCCGCTAACTCGAACGC 3’ and 5’ TTGTTTGGAGAGCCACCAGGC 3’), Serpine1 SE #2 (chr12:22634466-22634539, 5’ TTGAGTGGCAGACAGCCGACA 3’ and 5’ GGCGGCCTCCAACATTCCTC 3’), Serpine1 TSS (chr12:22640931-22641011, 5’ AGCCCCACCCACCTTCTAACTC 3’ and 5’ TACTGGGAGGGAGGGAAGGAGA 3’), Ctgf SE #1 (chr1:21871291-21871393, 5’ AGCCCTGGAATGCTGTTT 3’ and 5’ ACCGCATGATATCTCCTAAACC 3’), Ctgf SE #2 (chr1:21984665-21984753, 5’ AGTGAGTCAGGGAGGAAGAA 3’ and 5’ CTCCTGCAGCCTGTGATTAG 3’), Ctgf TSS (chr1:21854660-21854725, 5’ CAGACCCACTCCAGCTCCGA 3’ and 5’ GTGGCTCCTGGGGTTGTCCA 3’), and Fos TSS (chr6:109300463-109300526, 5’ GACTGGATAGAGCCGGCGGA 3’ and 5’ CAGAGCAGAGCTGGGTGGGA 3’).

### Protein extraction and western blot

Treated cardiomyocytes were lysed in RIPA buffer (1% NP-40, 50 mM Tris-HCl pH 7.4, 150 mM NaCl, 1 mM EDTA, 1 mM EGTA, 0.1% SDS, 0.5% sodium deoxycholate) and protein quantified by Bradford assay. Proteins were denatured at 65°C for 15 min in Laemmilli buffer and protein expression was assessed by western blot. Western blots were probed with anti-Brd4 (Bethyl, A301-985A; 1:1000) or anti-Hsp90 (Enzo Life Sciences, AC88; 1:1000) in 5% milk overnight at 4°C. The following day, blots were visualized with peroxidase-conjugated secondary antibodies and an Amersham^TM^ Imager 600.

### Statistical Analysis

All statistical analysis was performed using GraphPad Prism 8 software. Two-way analysis of variance was performed followed by post-hoc t-tests with Bonferroni correction (Figure 1B, Figure 2B/C/E, Figure 3B/C, Figure 5B/C, Figure 9A/C). Unpaired t-test was completed for validation of gene knockdown by siRNA (Figure 2D, Figure 5A) and for Brd4 ChIP with forskolin and IBMX (Figure 9B). One-way analysis of variance followed by Dunnett’s post-hoc comparison was performed for Brd4 ChIP (Figure 6A) and Brd4 protein expression (Figure 6C) following 24 h receptor activation. For the upstream regulator prediction (Figure 7G), the p-value was obtained by a Fisher’s Exact Test completed within the Ingenuity Pathway Analysis software.

## Acknowledgements

This work was supported by a grant from the Heart and Stroke Foundation of Canada (G-15-0008938) to T.E.H and J.C.T, a grant from Canadian Institute of Health Science (CIHR) (MOP 130-362) to J.C.T. and a grant from CIHR (PJT 159687) to T.E.H. R.M. was supported by a studentship from the McGill-CIHR Drug Development Training Program and McGill Faculty of Medicine. We thank Dr. Jin Zhang (UCSD), Dr. Karel Svoboda, and Novartis Institutes for Biomedical Research (Emeryville, CA) for providing materials instrumental to this study. We thank the McGill Imaging and Molecular Biology Platform for assistance with our high-content microscopy experiments. Lastly, we thank all the members of Dr. Hébert and Dr. Tanny’s labs for discussions and critical reading of the manuscript.

